# The Effects of N-linked Glycosylation on SLC6 Transporters

**DOI:** 10.1101/2022.07.12.499387

**Authors:** Matthew C. Chan, Diwakar Shukla

**Affiliations:** Department of Chemical and Biomolecular Engineering, University of Illinois Urbana-Champaign, Urbana, IL, 61801, United States; Center for Biophysics and Quantitative Biology, University of Illinois Urbana-Champaign, Urbana, IL, 61801, United States; Department of Plant Biology, University of Illinois Urbana-Champaign, Urbana, IL 61801, United States; Department of Bioengineering, University of Illinois Urbana-Champaign, Urbana, IL 61801, United States

## Abstract

Membrane transporters of the solute carrier 6 (SLC6) family mediate various physiological processes by facilitating the translocation of amino acids, neurotransmitters, and other metabolites. In the human body, these transporters are tightly controlled through various post-translational modifications with implications on protein expression, stability, membrane trafficking, and dynamics. While N-linked glycosylation is a universal regulatory mechanism among eukaryotes, the exact molecular mechanism of how glycosylation affects the SLC6 transporter family. It is generally believed that glycans influence transporter stability and membrane trafficking, however, the role of glycosylation on transporter dynamics remains inconsistent, with differing conclusions among individual transporters across the SLC6 family. In this study, we collected over 1 millisecond of aggregated all-atom molecular dynamics (MD) simulation data to identify the impact of N-glycans of four human SLC6 transporters: the serotonin transporter, dopamine transporter, glycine transporter, and neutral amino acid transporter B^0^AT1. We designed our computational study by first simulating all possible combination of a glycan attached to each glycosylation sites followed by investigating the effect of larger, oligo-N-linked glycans to each transporter. Our simulations reveal that glycosylation does not significantly affect transporter structure, but alters the dynamics of the glycosylated extracellular loop. The structural consequences of glycosylation on the loop dynamics are further emphasized in the presence of larger glycan molecules. However, no apparent trend in ligand stability or movement of gating helices was observed. In all, the simulations suggest that glycosylation does not consistently affect transporter structure and dynamics among the collective SLC6 family and should be characterized at a per-transporter level to further elucidate the underlining mechanisms of *in vivo* regulation.

## Introduction

The solute carrier 6 (SLC6) family is a class of secondary active co-transporters that mediates the reuptake of amino acids, biogenic amines, osmolytes, and metabolites, thereby maintaining cellular homeostasis throughout the body. ^1^ These transporters harness the energy of a favorable sodium ion concentration gradient to power the uphill transport of substrates across the plasma membrane. Many SLC6 transporters are also members of the neurotransmitter:sodium symporter (NSS) family and are essential for regulating neurotransmission in the central and peripheral nervous system.^2^

Members of the SLC6 family adopt the canonical 12 transmembrane (TM) helix LeuT fold with the transporter core formed by helices 1-5 and 6-10 arranged in a 5+5 inverted pseudo-symmetric repeat topology and two additional helices, 11 and 12, residing on the periphery of the core (Figure 1).^3^ The transport of substrates is dictated by the structural rearrangements that enables the transporter to alternate between an extracellular accessible or outward-facing (OF) conformation to the intracellular accessible or inward-facing (IF) conformation. Specifically, SLC6 transporters undergo a rocking-bundle mechanism in which the transmembrane helices 1 and 6 serve as gating helices that undergo a “rocking” conformational shift from the rigid scaffold domain, thus enabling the opening and closure of the orthosteric binding site.^4^ The recent determination of various SLC6 transporters and its bacterial homologs have established the structural basis of substrate and inhibitor molecule binding.^3,5–8^

**Figure 1:**
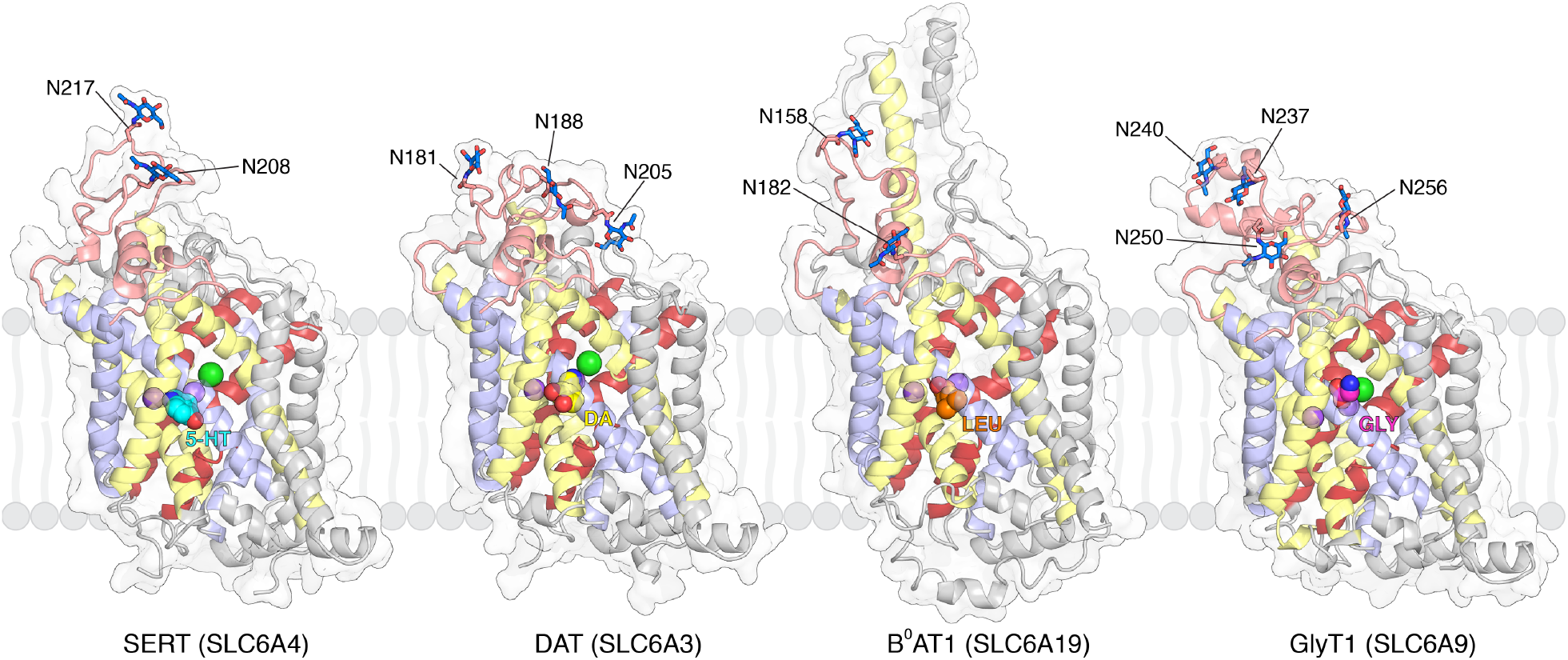
Starting structures of SLC6 transporters used for MD simulations. Transporters were modeled in the outward-facing conformation with substrates and ions initialed bound in the orthosteric pocket. Sodium and chloride ions are shown as purple and green spheres, respectively. Respective substrates are shown as spheres (5-HT: serotonin, DA: dopamine, LEU: leucine, GLY: glycine). The transporters are shown in cartoon representation and colored as follows: gating helices 1 and 6, red; 5+5 helix repeats, yellow and pale blue; extracellular loop 2, salmon. N-linked glycosylation sites with an N-acetylglucosamine glycan are represented as sticks and labeled accordingly.

Despite sharing 20-25% sequence identity with human SLC6 transporters, prokaryotic SLC6 proteins have historically illuminated the elusive structure-function relationships of this important class of transporters.^3,9–11^ While the general understanding of transport and conformational dynamics may be applied to characterize human transporters, the consequences of post-translational modifications cannot be inferred as prokaryotic homologs do not share the similar mechanisms or structural features of regulatory components as to their eukaryotic counterparts.^1,12^ As such, recent work has focused on elucidating the molecular mechanisms of post-translation modifications and its effect on human SLC6 transporters.^1,2^ These studies include phosphorylation,^13–17^ palmitoylation, ^18,19^ glycosylation,^20–24^ and ubiquitination^25^ and its implications on transporter dynamics, stability, oligomerization, trafficking, and uptake activity.

The glycosylation of SLC6 transporters has been widely documented to affect transporter activity;^20–24^ however, various mechanisms of how glycosylation mediates transporter function have been proposed for different SLC6 members.^2^ For example, glycosylation has been suggested to influence transporter stability in the membrane as demonstrated for the serotonin, dopamine, and norepinephrine transporters,^20,21,23^ whereas in glycine and GABA transporters, glycosylation regulates membrane trafficking.^22,24^ The removal of glycans did not affect ligand binding or transport function for the serotonin and norepinephrine transporters;^20,23^ however, mutagenesis of N-linked glycosylation sites in the dopamine and glycine 1 transporters resulted in reduced uptake rates. ^21,22^ Furthermore, the degree of glycosylation widely differs among expression organisms, tissues, and cell development,^26–28^ and as such, the extent of glycosylation and its effect on transporter structure and dynamics remain ambiguous.

With the surge in performance of graphical processing units and numerical algorithms, molecular dynamics (MD) simulations present a powerful approach to to characterize post-translational modifications and its effect on protein structure and dynamics. Recent applications of atomistic simulations to investigate post-translational modifications has identified how phosphorylation alters the hydrogen bonding network the serotonin transporter,^14^ glycosylation induces open conformations of the yeast disulfide isomerase,^29^ and nitration prevents ligand binding of a plant abscisic acid receptor.^30^ Moreover, MD simulations provide a technique to probe the structural dynamics in a label-free, fully atomistic approach, ideal for addressing the differences in experimental setup.

In this current work, we designed a computational study to systematically investigate the structural consequences of N-linked glycosylation on SLC6/NSS transporters. We performed microsecond MD simulations on four human SLC6/NSS transporters (Figure 1): the serotonin transporter (SERT, SLC6A4), the dopamine transporter (DAT, SLC6A3), the neutral amino acid transporter B^0^AT1 (SLC6A19), and the glycine transporter 1 (GlyT1, SLC6A9), to elucidate the role of glycans on transporter stability and conformational dynamics. We first examined the effects of glycosylation on the four transporters with glycans attached to each glycosylation site in a combinatorial fashion. In the second part of our study, we simulated the transporters with various degrees and complexity of olgioglycans to probe in the influence of larger glycan chains on the protein structure. Our simulations reveal that glycosylation does not significantly affect overall transporter structure, but alters the dynamics of the extracellular loops, but not in a sequence-dependent manner. Overall, we conclude that glycosylation does not significantly affect dynamics associated with substrate transport and thus is likely more involved in cellular sensing and regulation in the cell.

## Results

### Glycosylation does not significantly affect transporter structure but alters loop dynamics

The extracellular loop (EL) 2 of SLC6 transporters contain two to four N-linked glycosylation sites that follow the Asn-X-Ser/Thr amino acid sequence motif, where X is any residue except proline (Figure S1).^31^ Previous biophysical characterization of NSS transporters have reveal the extracellular loops to be coupled with the substrate transport dynamics,^32–34^ and as such, we hypothesized if the addition of bulky, hydrophobic glycans may affect the structure and dynamics of the transporter. We performed microsecond long MD simulations of four SLC6 transporters with a N-acetylglucosamine glycan modeled to each N-linked glycosylation site in a combinatorial fashion (Figure 1 and Table S1). Simulations were initiated from an outward-facing conformation with ions and respective substrates bound in the orthosteric binding site and embedded in a multicomponent phospholipid bilayer. A total of 29-30 MD replicates of 1*µ*s long simulations were collected for each transporter and glycosylation state, resulting in an aggregated simulation dataset of 949 *µ*s (Table S1).

The root-mean-square deviation (RMSD) and fluctuations (RMSF) with respect to the initial starting structure is presented in Figure 2. The simulations reveal that glycosylation does not significantly affect the overall transporter structure, with the exception of SERT-N208 and DAT-N181-N188 exhibited a marginal decrease in RMSD as compared to the deglycosylated transporter (Figure 2A). When specifically examining the structure of EL2 alone, the simulations of glycosylated B^0^AT1 and GlyT1 were not observed to significantly differ from the respective deglycosylated transporters (Figure 2B). In contrast, two glycosylated SERT systems and three glycosylated DAT systems were found to have a significant decrease in EL2 RMSD (Figure 2B). The averaged RMSF of EL2 further reveals the same two glycosylated SERT systems (N208, N208-N217) to experience decreased dynamics (Figure 2C). However, for the other studied transporters, only a doubly glycosylated DAT (N188-N205) and a triply glycosylated GlyT1 (N237-N240-N250) were found to have significantly decreased EL2 fluctuations. In the remaining transporters, including all the B^0^AT1 simulations, glycosylation was not observe to profound impact on EL2 dynamics.

**Figure 2:**
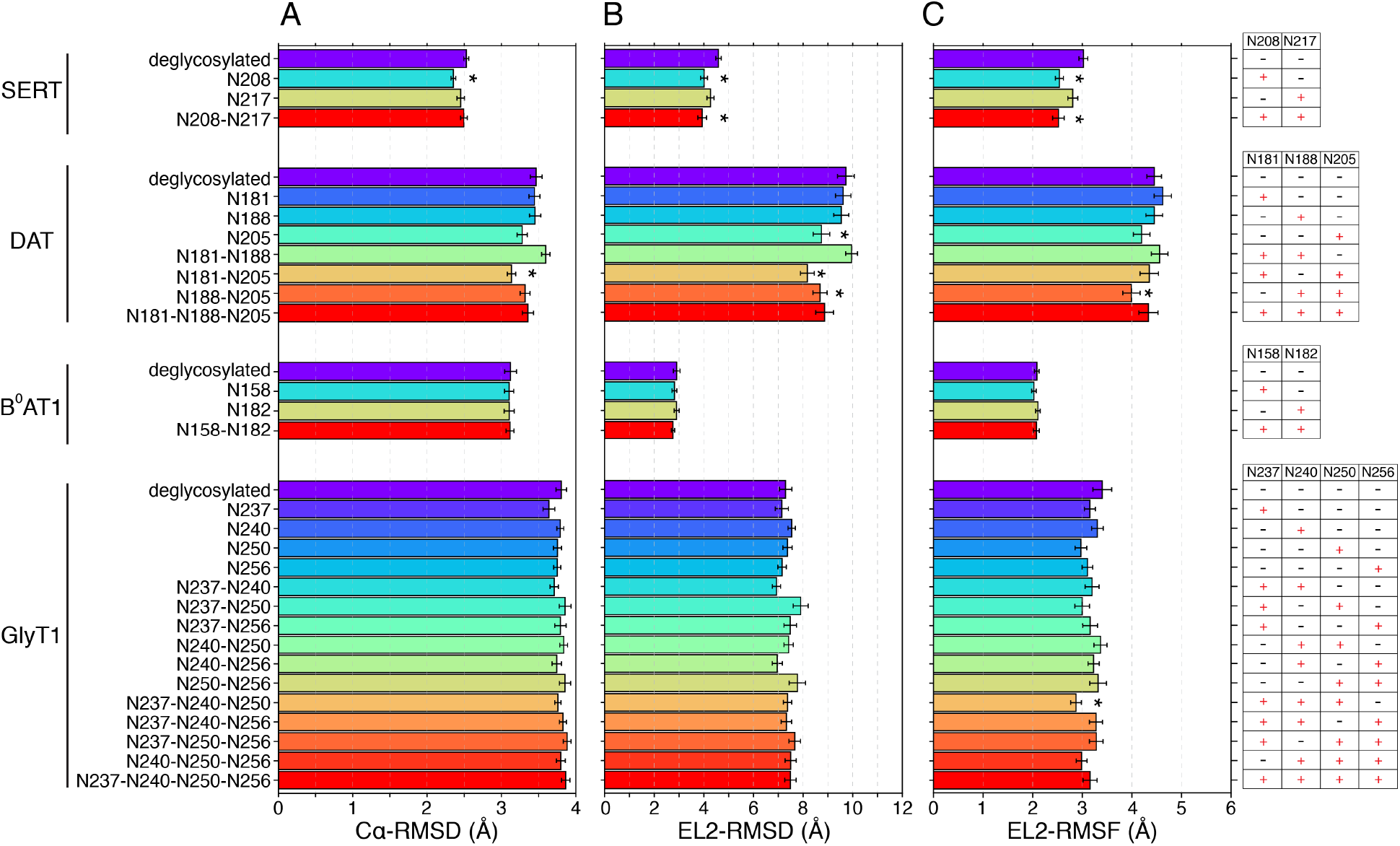
Glycosylation does not significantly affect transporter structure. Structural measurments of **(A)** the root-mean-square deviations (RMSD) of all transporter C*α* atoms, **(B)** RMSD of extracellular loop 2 (EL2), and **(C)** averaged root-mean-square fluctuations (RMSF) of extracellular loop 2 atomic displacement. The initial structure used for MD simulations was used as the respective reference for all calculations. Quantities are averaged from all 1*µ*s MD replicates for each respective system. Error bars represent standard error across the replicates. * indicates values significantly different (*p*-value < 0.05, independent *t*-test) from the respective deglycosylated transporter. A table depicting the N-glycosylated residues is shown on the right, with (+) indicating a N-acetylglucosamine glycan was added and (-) as deglycosylated.

Figure 3 shows the difference per-residue RMSF with respected to the deglycosylated transporter for the four studied SLC6 transporters. The plots reveal that glycosylation alters the dynamics of EL2 in a differing manner among transporters (Figure 3). In SERT simulations, glycosylation consistently decreases the fluctuations of EL2 as compared to the deglycosyated SERT (Figure 3A). However, the effects of glycosylation on EL2 dynamics varies and does not show a consistent trend among DAT and GlyT1 transporters (Figure 3B, D). In DAT specifically, we observed the fully glycosylated transporter (N181-N188-N205) to increase the dynamics of the cytoplasmic base of TM5 (Figure 3B). Extensive literature supports the unwinding of the TM5 as a key structural rearrangement for propagating transition to the inward-facing state.^32,34–36^ Furthermore, the number of glycosylated Asn residues did not appear to be correlated with effects on dynamics. Interestingly in B^0^AT1, glycosylation did not have a pronounced effect on EL2 dynamics, but allosterically alters the displacements of the nearby EL4 (Figure 3C). The glycans were not observed to come into contact with EL4, but the cryo-EM complex reveals that EL4 and the extended TM7 play a role in trafficking and interfacing with the angiotensin-converting enzyme 2.^37^ EL4 of B^0^AT1 contains a number of N-linked glycosylation sites,^37^ but the effects of these sites were not investigated in this study.

**Figure 3:**
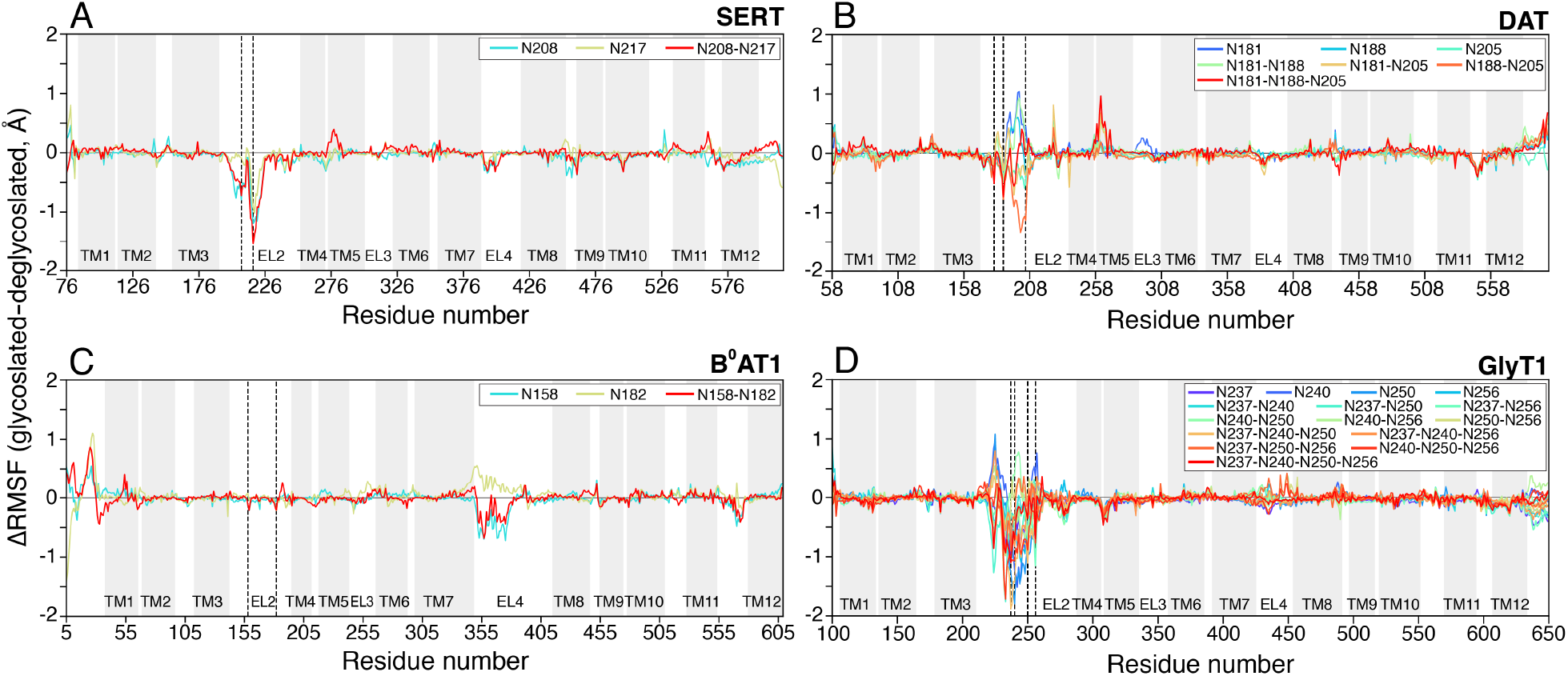
Difference RMSF plots of the glycosylated transporters. The difference RMSF (ΔRMSF) in which the RMSF of the deglycosylated transporter was subtracted from the glycosylated transporter is plotted along the primary residue sequence. The initial structure used for MD simulations was used as the respective reference for RMSF calculations. Quantities are averaged from all 1*µ*s MD replicates for each respective system. The glycosylated systems are plotted and colored according to Figure 2. Transmembrane helices are marked in gray regions along the residue numbers. N-linked glycosylation sites are marked in black dashed line.

Overall, though the difference in deviations is of relatively small magnitude (∼0.5-1.5 Å), N-glycosylation of EL2 does not consistently affect the structural dynamics within individual transporters and across the sampled SLC6 family. From the simulations, we observed marginal differences (< 1Å) in the distance distributions of gating helices, thus suggesting that N-glycosylation does not have a profound effect on transport dynamics (Figure S2). Furthermore, glycosylation does not consistently affect the stability of the ligand bound in the orthosteric site, with the exception of SERT (Figure S3). The simulations of SERT and its glycosylated forms reveal that the RMSD of the serotonin, with respect to the initial bound pose, is decreased thus suggesting greater ligand stability upon glycosylation (Figure S3A). In all, the simulations reveal that the presence of hydrophobic glycans on a solvent-exposed domain of the transporter alters its local environment, but does not extend to the remainder of the transporter.

### Oligo-N-linked glycosylation further stabilizes loop fluctuations

Though ubiquitous among eukaryotes, it is evident to note that the degrees of glycosylation and its regulatory role widely differs among species and cell types.^28^ The previous body of literature has extensive explored the use of cell lines from a variety of organisms including, but not limited to, human (HEK-293), ^21^ insect (Sf9),^20^ monkey (COS),^22^ pig (LLC-PK1),^23^ and hamster (CHO).^24^ Moreover, in humans, N-glycosylation patterns have been noted to differ among various cell types and developmental stages^26,27^ and thus further illuminates the complexity of glycosylation in the nervous system and throughout the body.^38^

As it is not feasible to investigate all possible glycan and linkage patterns, nor has it been characterized in exact detail, we designed MD systems of the four studied transporters in a pattern of increasing glycans moieties to serve as a representative and general model of complex oligoglyans (Figure 4A). The complex glycans ranged from a linear 2 N-acetylglucosamine glycans to a branch olgioglycan containing 9 carbohydrates in total. For simulations of the oligoglycans, all Asn glycosylation sites on EL2 were modeled as in the glycosylated form. Figure 4B shows a representative structure of 9-glycan system for SERT. The glycosylated transporters were constructed in the same protocol as the single N-acetylglucosamine glycosylated transporter systems and a total of 29-30 replicates were simulated for 1*µ*s each, totaling in an additional 359 *µ*s of aggregate data.

**Figure 4:**
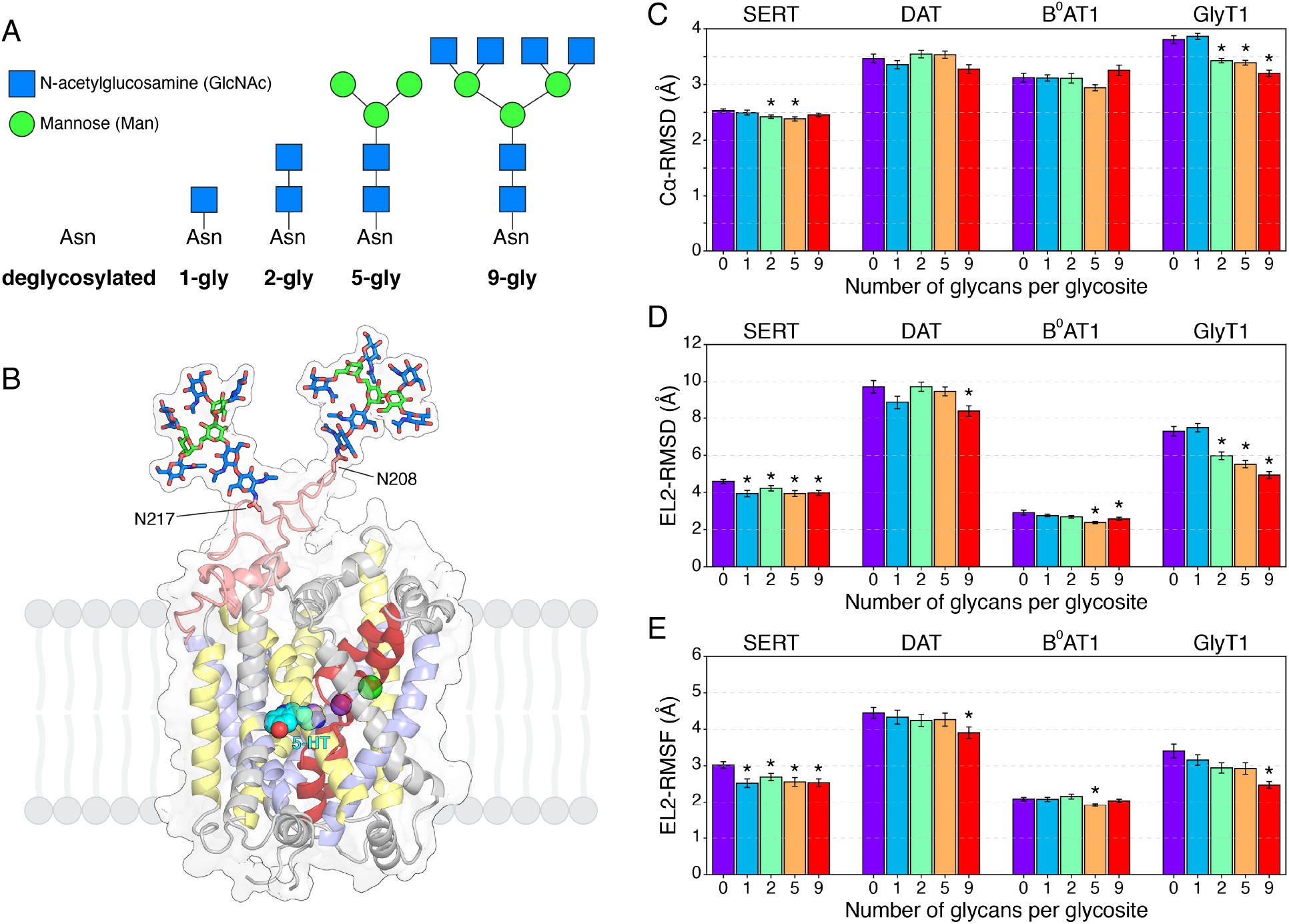
Structural effects of oligo-N-linked glycans. **(A)** The oligo-N-linked glycans modeled in this study. The oligoglycans were added to all glycoslation sites for each transporter and simulated under the same protocol as the single N-acetylglucosamine glycan simulations. **(B)** Representative structure of SERT and the 9-glycan group added to both glycosylation sites. Transmembrane helices and substrates are represneted and colored according to Figure 1. **(C, D, E)** Structural measurements of the oligoglycan-transporter systems, similarly calculated as to Figure 2. The values of **(C)** C*α* RMSD, **(D)** EL2 RMSD, and **(E)**, averaged EL2 RMSF were averaged from the 1*µ*s MD replicates for each respective system. Error bars represent standard error across the replicates. * indicates values significantly different (*p*-value < 0.05, independent *t*-test) from the respective deglycosylated (0 glycan) transporter.

Similar to simulations of the single N-acetylglucosamine glycosylated transporters, the simulations of the complex oligoglycans further suggest that N-glycosylation does not uniformly affect SLC6 transporters. With regards to the overall transporter structure, both DAT and B^0^AT1 when glycosylated to any degree were not observed to display differences among the sets of simulations (Figure 4C). The glycosylated SERT systems shown minimal differences in overall transporter RMSD (< 0.5 Å), though the 2- and 5-glycan systems were indicated as significant when compared to deglycosylated SERT. When examining the structure and dynamics of EL2, the simulations for the largest simulated glycans reveal that the RMSD of DAT and B^0^AT1 EL2 to be significantly lower compared to the deglycosylated transporter (Figure 4D). In SERT, the single N-acetylglucosamine added to both glycosylation site N208 and N217 was observed to decrease the EL2 RMSD (Figure 2B) and the presence of larger glycans did not significantly further influence the EL2 structure (Figure 4D, E). Most strikingly, increasing the number of glycans and complexity of linkages added to GlyT1 was correlated with a decrease in overall transporter and EL2 RMSD (Figure 4C, D).

The simulations further reveal that the dynamics of EL2 are reduced when glycosylated with complex oligoglycans, with the most significant differences observed with more glycans added per glycosylation site (Figure 4E). The RMSF plots of the olgio-glycosylated transporters illustrate that the fluctuations of EL2 are also altered compared to the deglycosylated transporter (Figure 5). The dynamics of EL2 on SERT and B^0^AT1 do not differ widely from the single N-acetylglucosamine glycosyated transporters (Figure 5A, C). However, more notably, the fluctuations for DAT and GlyT1 EL2 when glycosylated with the oligoglycans are generally diminished across all degrees of glycosylation (Figure 5B, D). Furthermore, the presence of the complex oligoglycans did not consistently affect the motions of the gating helices (Figure S4) nor the stability of the bound ligand as similarly observed in the single N-acetylglucosamine transporter simulations (Figure S5).

**Figure 5:**
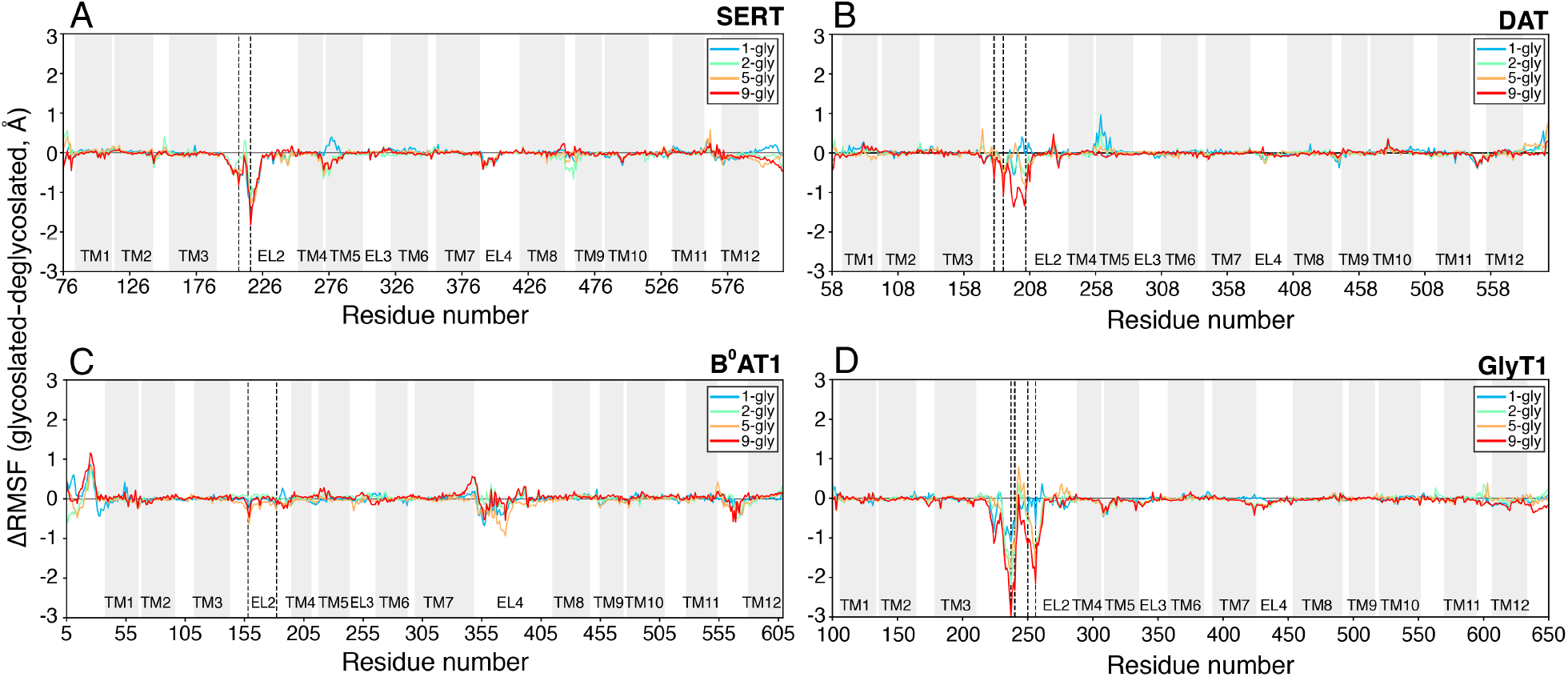
Difference RMSF plots of the oligo-glycosylated transporters. The difference RMSF (ΔRMSF) in which the RMSF of the deglycosylated (0-gly) transporter was subtracted from the glycosylated transporter is plotted along the primary residue sequence. The initial structure used for MD simulations was used as the respective reference for RMSF calculations. Quantities are averaged from all 1*µ*s MD replicates for each respective system. MD systems with varying degrees of glycosylation are plotted and colored according to Figure 4. Transmembrane helices are marked in gray regions along the residue numbers. N-linked glycosylation sites are marked in black dashed line.

## Discussion

The activity of SLC6 transporters are tightly controlled through intricate regulatory mechanisms. Consequently, dysregulation of transport activity is associated with various neurodegenerative, respiratory, and cardiovascular diseases.^1,2^ As these transporters are essential for maintaining cellular homeostasis, understanding the conformational heterogeneity and how post-translational modification alter the underlying dynamics is pivotal for designing effective therapeutic molecules.

In this study, we investigated the structural effects of N-linked glycosylation on four human SLC6 transporters using MD simulations in two sets of computational experiments. In first simulating all possible combinations of N-linked glycosylation on the EL2 of the studied transporters, we observed a few significant differences in overall transporter RMSD and EL2 dynamics as compared to the deglycosylated transporter. The RMSF plots further show that glycosylation reduces the dynamics of EL2 of SERT, is indiscernible from the deglycosylated system in B^0^AT1, and ununiformly affects DAT and GlyT1 EL2. However, in the second set of simulations, when complex olgioglycans are attached to the transporters, the EL2 dynamics were generally decreased. In both sets of simulations, single and oligoglycan, we did not observe discernible differences in the dynamics of the gating helices nor the stability of the bound ligand within the simulated timescales. As such, we conclude that the simulations support the existing literature that glycosylation does not have significant effect on the substrate transport dynamics and is likely more involved in maintaining stability and proper trafficking in the membrane.

Glycosylation is an essential and universal post-translation modification for regulating protein function. The use of MD simulations enables an atomistic characterization of the structural and dynamic consequences of glycosylation^29,39,40^ and other post-translational modifications. ^14,41,42^ Though, glycosylation has been widely understood to affect SLC6 transporter stability and trafficking,^1,2^ our simulations show that N-glycosylation minimally affects overall transporter dynamics, but reduces the fluctuation of the extracellular loops. However, we did not observe glycosylation to consistently alter SLC6 transporter structure which may further explain the differences in regulatory function previously characterized experimentally.^20–24^ Furthermore, previous simulations of glycoproteins further underlines a lack of uniformity in regulating protein structure and dynamics^39,43,44^ and may suggest that the disruption of the local protein environment has a greater role in modulating dynamics and stability rather than the glycans itself.

## Methods

### System preparation

To investigate the effects of glycosylation on transporter dynamics and stability, we selected four human transporters from the SLC6/NSS family: the serotonin transporter (SERT), dopamine transporter (DAT), neutral amino-acid transporter B^0^AT1, and the glycine transporter 1 (GlyT1). These transporters have extensive structural and/or biochemical characterization of the effects of glycosylation.^5–7,20–22,37^

We initiated all simulations from an outward-facing conformation with the transporter’s respective substrates bound in the orthosteric pocket. The initial structures were obtained as followed: SERT, three-dimensional coordinates from the outward-facing crystal structure (PDB: 5IX6) with with Na1, Na2, Cl^-^ bound and serotonin (5-HT) modeled based on our previous MD simulation study;^32^ DAT, a homology model based on the outward-facing *Drosophila melanogaster* DAT crystal structure (PDB: 4XP1) with Na1, Na2, Cl^-^ and dopamine (DA) modeled based on the crystal structure; B^0^AT1: three-dimensional coordinates from the outward-facing cryo-EM structure (PDB: 6M18) with with Na1, Na2, and leucine based on the structural alignment with Leu-bound LeuT (PDB: 2A65); and GlyT1, a homology model based on the outward-facing *Drosophila melanogaster* DAT crystal structure previously modeled by Zhang *et al*.^45^ with Na1, Na2, Cl^-^ bound and glycine bound based on the structural alignment with Gly-bound LeuT (PDB: 3F4J). The GlyT1 model did not initially contain extracellular loop 2 (EL2) and as such, we modeled the loop using the comparative modeling module of the ROBETTA web server.^46^ The resulting EL2 model displayed alpha helical secondary structure elements at residues 235 to 239 and 243 to 252, which is further suggested by the IUPRED intrinsic disorder structure prediction web server.^47^

The transporters were embedded in a 90 × 90 Å^2^ multi-component phospholipid bilayer using the CHARMM-GUI web server.^48^ For SERT, DAT, and GlyT1, the transporter was embedded in a 2:1 POPC:POPE symmetric lipid bilayer, loosely based on the neuronal plasma membrane composition.^49^ As B^0^AT1 is expressed in the membrane of the small intestine,^50^ we embedded the transporter in a 3:2:1 POPE:POPC:POPS membrane to mimic its native environment.^51^ We note the exclusion of cholesterol molecules in the simulated membranes. While cholesterol is physiological relevant in the human membrane environment, it has been extensively shown to sterically stabilize outward-facing conformations.^52,53^ As such, we excluded cholesterol to prevent unintended inhibition of transporter dynamics. N- and C-termini were capped with acetyl and methyl amide groups, respectively. Titratable residues were modeled in accordance to pK_a_ calculations using PROPKA3.0.^54^ The systems were solvated with TIP3P water molecules and 150 mM NaCl. The mass of hydrogen atoms and connecting atoms were repartitioned accordingly to Hopkins *et. al*.^55^ For single-glycan simulations, an N-acetylglucosamine glycan was modeled to Asn glycosylation sites in a combinatorial fashion. For simulations of olgioglycans (2-gly, 5-gly, and 9-gly), the glycans were simultaneously modeled on all Asn glycosylation sites. Individual details of constructed system are presented in Table S1 and S2.

### Molecular dynamics simulations

Prior to production, the systems were minimized and equilibrated using the AMBER18 MD package employing CHARM36m force field. The CHARMM *psf* topology and coordinate files were converted to AMBER *prmtop* and *rst7* file using the *chamber* module of the ParmED package.^56^ Each system was first subjected to an energy minimization protocol of 5,000 steps using the steepest descent method, followed by 45,000 minimization steps using the conjugate gradient method. The systems were then heated to 300K for 5 ns in a constant particle, pressure, temperature (NPT) ensemble while the protein backbone, bound substrates, and glycans were restrained with a force constant of 5 kcal/mol-Å^2^. The equlibrated snapshot was then converted to an OpenMM system parameterized with an OpenMM ForceField using the CHARM36m force field.^57^

Production simulations were performed using the OpenMM 7.7.0 package^58^ on either the Folding@Home distributing computing platform^59^ or the University of Illinois National Center for Supercomputing Applications Delta supercomputer. Langevin dynamics was performed using a Langevin integrator using an integration timestep of 4 fs, temperature of 300 K, and collision rate of 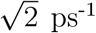. The system pressure of 1 bar was maintained using the Monte Carlo Membrane Barostat with a surface tension of 200 bar-nm and update frequency of 100 steps. Nonbonded forces were calculated using the particle mesh Ewald method with a 12 Å cutoff distance. Simulations were performed using mixed numerical precision, periodic boundary conditions, and hydrogen mass repartitioning.^55^ A total of 30 MD replicates for each system with different initial velocities were sent to Folding@Home users and simulated up to 1 *µ*s. Trajectories in which simulation data was not received from Folding@Home clients were not used for analysis. In all, a total of 29-30 1*µ*s long trajectories for each glycosylated system and transporter were collected and analyzed (Table S1 and S2). Trajectory snapshots were saved every 100 ps during production simulations.

### Trajectory analysis

Trajectories were processed with in-house scripts utilizing CPPTRAJ, pytraj, and MDTraj packages^60,61^ and visualized using Visual Molecular Dynamics (VMD) ^62^ and PyMOL. The root-mean-square deviation (RMSD) of atomic positions were calculated on only C*α* atoms. The root-mean-square fluctuations (RMSF) of each residue was calculated on all atoms and mass-averaged by residue. The initial structure used for production simulations was used as the reference for these calculations. An independent *t*-test was performed to compare RMSD and RMSF of the glycosylated and the respective deglycosylated system with a significance level of 0.05. C*α* atoms used for the center-of-mass calculations for the distance distribution of the gating helices are listed in Table S3. Plots were generated using the matplotlib Python library.

## Supporting information

Supplementary Methods, Images, Tables and Results

## Acknowledgements

This work was supported by an NSF Early Career Award by NSF MCB 18-45606 to D.S. This research is also part of the Delta research computing project, which is supported by the National Science Foundation (award OCI 2005572), and the State of Illinois. Delta is a joint effort of the University of Illinois Urbana-Champaign and its National Center for Supercomputing Applications. The authors thank Folding@Home donors for computational resources for this project. M.C.C. thanks Austin T. Weigle for insightful discussion and Nicole Chiang for technical assistance pertaining to this study.

## Author contributions statement

D.S. acquired funding for this project. D.S. and M.C.C. designed the study. M.C.C performed the simulations. M.C.C. and D.S. analyzed the results. M.C.C. and D.S. prepared the manuscript.

## References

(1) Pramod, A. B.; Foster, J.; Carvelli, L.; Henry, L. K. SLC6 transporters: Structure, function, regulation, disease association and therapeutics. Molecular Aspects of Medicine 2013, 34, 197–219.

(2) Kristensen, A. S.; Andersen, J.; Jørgensen, T. N.; Sørensen, L.; Eriksen, J.; Loland, C. J.; Strømgaard, K.; Gether, U. SLC6 Neurotransmitter Transporters: Structure, Function, and Regulation. Pharmacological Reviews 2011, 63, 585–640.

(3) Yamashita, A.; Singh, S. K.; Kawate, T.; Jin, Y.; Gouaux, E. Crystal structure of a bacterial homologue of Na+/Cl–dependent neurotransmitter transporters. Nature 2005, 437, 215–223.

(4) Forrest, L. R.; Rudnick, G. The Rocking Bundle: A Mechanism for Ion-Coupled Solute Flux by Symmetrical Transporters. Physiology 2009, 24, 377–386.

(5) Penmatsa, A.; Wang, K. H.; Gouaux, E. X-ray structure of dopamine transporter elucidates antidepressant mechanism. Nature 2013, 503, 85–90.

(6) Coleman, J. A.; Green, E. M.; Gouaux, E. X-ray structures and mechanism of the human serotonin transporter. Nature 2016, 532, 334–339.

(7) Shahsavar, A.; Stohler, P.; Bourenkov, G.; Zimmermann, I.; Siegrist, M.; Guba, W.; Pinard, E.; Sinning, S.; Seeger, M. A.; Schneider, T. R.; Dawson, R. J. P.; Nissen, P. Structural insights into the inhibition of glycine reuptake. Nature 2021, 591, 677–681.

(8) Motiwala, Z.; Aduri, N. G.; Shaye, H.; Han, G. W.; Lam, J. H.; Katritch, V.; Cherezov, V.; Gati, C. Structural basis of GABA reuptake inhibition. Nature 2022, In press.

(9) Quick, M.; Yano, H.; Goldberg, N. R.; Duan, L.; Beuming, T.; Shi, L.; Weinstein, H.; Javitch, J. A. State-dependent Conformations of the Translocation Pathway in the Ty-(1) rosine Transporter Tyt1, a Novel Neurotransmitter:Sodium Symporter from Fusobacterium nucleatum. Journal of Biological Chemistry 2006, 281, 26444–26454.

(10) Androutsellis-Theotokis, A.; Goldberg, N. R.; Ueda, K.; Beppu, T.; Beckman, M. L.; Das, S.; Javitch, J. A.; Rudnick, G. Characterization of a Functional Bacterial Homologue of Sodium-dependent Neurotransmitter Transporters. Journal of Biological Chemistry 2003, 278, 12703–12709.

(11) Quick, M.; Javitch, J. A. Monitoring the function of membrane transport proteins in detergent-solubilized form. Proceedings of the National Academy of Sciences 2007, 104, 3603–3608.

(12) Macek, B.; Forchhammer, K.; Hardouin, J.; Weber-Ban, E.; Grangeasse, C.; Mijakovic, I. Protein post-translational modifications in bacteria. Nature Reviews Microbiology 2019, 17, 651–664.

(13) Quinlan, M. A.; Krout, D.; Katamish, R. M.; Robson, M. J.; Nettesheim, C.; Gresch, P. J.; Mash, D. C.; Henry, L. K.; Blakely, R. D. Human Serotonin Transporter Coding Variation Establishes Conformational Bias with Functional Consequences. ACS Chemical Neuroscience 2019, 10, 3249–3260.

(14) Chan, M. C.; Procko, E.; Shukla, D. Structural Rearrangement of the Serotonin Transporter Intracellular Gate Induced by Thr276 Phosphorylation. ACS Chemical Neuroscience 2022, 13, 933–945.

(15) Khoshbouei, H.; Sen, N.; Guptaroy, B.; Johnson, L.; Lund, D.; Gnegy, M. E.; Galli, A.; Javitch, J. A. N-Terminal Phosphorylation of the Dopamine Transporter Is Required for Amphetamine-Induced Efflux. PLoS Biology 2004, 2, e78.

(16) Vargas-Medrano, J.; Castrejon-Tellez, V.; Plenge, F.; Ramirez, I.; Miranda, M. PKCβ-dependent phosphorylation of the glycine transporter 1. Neurochemistry International 2011, 59, 1123–1132.

(17) Cristóvão-Ferreira, S.; Vaz, S. H.; Ribeiro, J. A.; Sebastião, A. M. Adenosine A2A receptors enhance GABA transport into nerve terminals by restraining PKC inhibition of GAT-1. Journal of Neurochemistry 2009, 109, 336–347.

(18) Foster, J. D.; Vaughan, R. A. Palmitoylation Controls Dopamine Transporter Kinetics, Degradation, and Protein Kinase C-dependent Regulation. Journal of Biological Chemistry 2011, 286, 5175–5186.

(19) Zeppelin, T.; Pedersen, K. B.; Berglund, N. A.; Periole, X.; Schiøtt, B. Effect of palmitoylation on the dimer formation of the human dopamine transporter. Scientific Reports 2021, 11.

(20) Tate, C. G.; Blakely, R. D. The effect of N-linked glycosylation on activity of the Na(+)- and Cl(-)-dependent serotonin transporter expressed using recombinant baculovirus in insect cells. Journal of Biological Chemistry 1994, 269, 26303–26310.

(21) Li, L.-B.; Chen, N.; Ramamoorthy, S.; Chi, L.; Cui, X.-N.; Wang, L. C.; Reith, M. E. The Role of N-Glycosylation in Function and Surface Trafficking of the Human Dopamine Transporter. Journal of Biological Chemistry 2004, 279, 21012–21020.

(22) Olivares, L.; Aragón, C.; Giménez, C.; Zafra, F. The Role of N-Glycosylation in the Targeting and Activity of the GLYT1 Glycine Transporter. Journal of Biological Chemistry 1995, 270, 9437–9442.

(23) Melikian, H. E.; Ramamoorthy, S.; Tate, C. G.; Blakely, R. D. Inability to N-glycosylate the human norepinephrine transporter reduces protein stability, surface trafficking, and transport activity but not ligand recognition. Molecular Pharmacology 1996, 50, 266–276.

(24) Cai, G.; Salonikidis, P. S.; Fei, J.; Schwarz, W.; Schülein, R.; Reutter, W.; Fan, H. The role of N-glycosylation in the stability, trafficking and GABA-uptake of GABA-transporter 1. FEBS Journal 2005, 272, 1625–1638.

(25) Miranda, M.; Sorkin, A. Regulation of Receptors and Transporters by Ubiquitination: New Insights into Surprisingly Similar Mechanisms. Molecular Interventions 2007, 7, 157–167.

(26) Lew, R.; Vaughan, R.; Simantov, R.; Wilson, A.; Kuhar, M. J. Dopamine transporters in the nucleus accumbens and the striatum have different apparent molecular weights. Synapse 1991, 8, 152–153.

(27) Patel, A. P.; Cerruti, C.; Vaughan, R. A.; Kuhar, M. J. Developmentally regulated glycosylation of dopamine transporter. Developmental Brain Research 1994, 83, 53–58.

(28) Croset, A.; Delafosse, L.; Gaudry, J.-P.; Arod, C.; Glez, L.; Losberger, C.; Begue, D.; Krstanovic, A.; Robert, F.; Vilbois, F.; Chevalet, L.; Antonsson, B. Differences in the glycosylation of recombinant proteins expressed in HEK and CHO cells. Journal of Biotechnology 2012, 161, 336–348.

(29) Weiß, R. G.; Losfeld, M.-E.; Aebi, M.; Riniker, S. N-Glycosylation Enhances Conformational Flexibility of Protein Disulfide Isomerase Revealed by Microsecond Molecular Dynamics and Markov State Modeling. The Journal of Physical Chemistry B 2021, 125, 9467–9479.

(30) Shukla, S.; Zhao, C.; Shukla, D. Dewetting controls plant hormone perception and initiation of drought resistance signaling. Structure 2019, 27, 692–702.

(31) Helenius, A.; Aebi, M. Roles of N-Linked Glycans in the Endoplasmic Reticulum. Annual Review of Biochemistry 2004, 73, 1019–1049.

(32) Chan, M. C.; Selvam, B.; Young, H. J.; Procko, E.; Shukla, D. The substrate import mechanism of the human serotonin transporter. Biophysical Journal 2022, 121, 715–730.

(33) Nielsen, A. K.; Möller, I. R.; Wang, Y.; Rasmussen, S. G. F.; Lindorff-Larsen, K.; Rand, K. D.; Loland, C. J. Substrate-induced conformational dynamics of the dopamine transporter. Nature Communications 2019, 10, 2714.

(34) Merkle, P. S.; Gotfryd, K.; Cuendet, M. A.; Leth-Espensen, K. Z.; Gether, U.; Loland, C. J.; Rand, K. D. Substrate-modulated unwinding of transmembrane helices in the NSS transporter LeuT. Science Advances 2018, 4.

(35) Malinauskaite, L.; Quick, M.; Reinhard, L.; Lyons, J. A.; Yano, H.; Javitch, J. A.; Nissen, P. A mechanism for intracellular release of Na+ by neurotransmitter/sodium symporters. Nature Structural & Molecular Biology 2014, 21, 1006–1012.

(36) Zhang, Y.-W.; Rudnick, G. The Cytoplasmic Substrate Permeation Pathway of Serotonin Transporter. Journal of Biological Chemistry 2006, 281, 36213–36220.

(37) Yan, R.; Zhang, Y.; Li, Y.; Xia, L.; Guo, Y.; Zhou, Q. Structural basis for the recognition of SARS-CoV-2 by full-length human ACE2. Science 2020, 367, 1444–1448.

(38) Scott, H.; Panin, V. M. Advances in Neurobiology; Springer New York, 2014; pp 367–394.

(39) Lee, H. S.; Qi, Y.; Im, W. Effects of N-glycosylation on protein conformation and dynamics: Protein Data Bank analysis and molecular dynamics simulation study. Scientific Reports 2015, 5.

(40) Shental-Bechor, D.; Levy, Y. Effect of glycosylation on protein folding: A close look at thermodynamic stabilization. Proceedings of the National Academy of Sciences 2008, 105, 8256–8261.

(41) Kuzmanic, A.; Sutto, L.; Saladino, G.; Nebreda, A. R.; Gervasio, F. L.; Orozco, M. Changes in the free-energy landscape of p38α MAP kinase through its canonical activation and binding events as studied by enhanced molecular dynamics simulations. eLife 2017, 6, e22175.

(42) Moffett, A. S.; Bender, K. W.; Huber, S. C.; Shukla, D. Allosteric Control of a Plant Receptor Kinase through S-Glutathionylation. Biophysical Journal 2017, 113, 2354–2363.

(43) Price, J. L.; Shental-Bechor, D.; Dhar, A.; Turner, M. J.; Powers, E. T.; Gruebele, M.; Levy, Y.; Kelly, J. W. Context-Dependent Effects of Asparagine Glycosylation on Pin WW Folding Kinetics and Thermodynamics. Journal of the American Chemical Society 2010, 132, 15359–15367.

(44) Fonseca-Maldonado, R.; Vieira, D. S.; Alponti, J. S.; Bonneil, E.; Thibault, P.; Ward, R. J. Engineering the Pattern of Protein Glycosylation Modulates the Thermostability of a GH11 Xylanase. Journal of Biological Chemistry 2013, 288, 25522–25534.

(45) Zhang, Y.-W.; Uchendu, S.; Leone, V.; Bradshaw, R. T.; Sangwa, N.; Forrest, L. R.; Rudnick, G. Chloride-dependent conformational changes in the GlyT1 glycine transporter. Proceedings of the National Academy of Sciences 2021, 118, e2017431118.

(46) Kim, D. E.; Chivian, D.; Baker, D. Protein structure prediction and analysis using the Robetta server. Nucleic Acids Research 2004, 32, W526–W531.

(47) Mészáros, B.; Erdős, G.; Dosztányi, Z. IUPred2A: context-dependent prediction of protein disorder as a function of redox state and protein binding. Nucleic Acids Research 2018, 46, W329–W337.

(48) Jo, S.; Kim, T.; Iyer, V. G.; Im, W. CHARMM-GUI: A web-based graphical user interface for CHARMM. Journal of Computational Chemistry 2008, 29, 1859–1865.

(49) Wilson, K. A.; MacDermott-Opeskin, H. I.; Riley, E.; Lin, Y.; O’Mara, M. L. Understanding the Link between Lipid Diversity and the Biophysical Properties of the Neuronal Plasma Membrane. Biochemistry 2020, 59, 3010–3018.

(50) dit Bille, R. N. V. et al. Human intestine luminal ACE2 and amino acid transporter expression increased by ACE-inhibitors. Amino Acids 2014, 47, 693–705.

(51) Forstner, G. G.; Tanaka, K.; Isselbacher, K. J. Lipid composition of the isolated rat intestinal microvillus membrane. Biochemical Journal 1968, 109, 51–59.

(52) Laursen, L.; Severinsen, K.; Kristensen, K. B.; Periole, X.; Overby, M.; Müller, H. K.; Schiøtt, B.; Sinning, S. Cholesterol binding to a conserved site modulates the conformation, pharmacology, and transport kinetics of the human serotonin transporter. Journal of Biological Chemistry 2018, 293, 3510–3523.

(53) Zeppelin, T.; Ladefoged, L. K.; Sinning, S.; Periole, X.; Schiøtt, B. A direct interaction of cholesterol with the dopamine transporter prevents its out-to-inward transition. PLOS Computational Biology 2018, 14, e1005907.

(54) Olsson, M. H. M.; Søndergaard, C. R.; Rostkowski, M.; Jensen, J. H. PROPKA3: Consistent Treatment of Internal and Surface Residues in Empirical pKa Predictions. Journal of Chemical Theory and Computation 2011, 7, 525–537.

(55) Hopkins, C. W.; Le Grand, S.; Walker, R. C.; Roitberg, A. E. Long-time-step molecular dynamics through hydrogen mass repartitioning. Journal of Chemical Theory and Computation 2015, 11, 1864–1874.

(56) Shirts, M. R.; Klein, C.; Swails, J. M.; Yin, J.; Gilson, M. K.; Mobley, D. L.; Case, D. A.; Zhong, E. D. Lessons learned from comparing molecular dynamics engines on the SAMPL5 dataset. Journal of Computer-Aided Molecular Design 2016, 31, 147–161.

(57) Huang, J.; Rauscher, S.; Nawrocki, G.; Ran, T.; Feig, M.; de Groot, B. L.; Grubmüller, H.; MacKerell, A. D. CHARMM36m: an improved force field for folded and intrinsically disordered proteins. Nature Methods 2016, 14, 71–73.

(58) Eastman, P.; Swails, J.; Chodera, J. D.; McGibbon, R. T.; Zhao, Y.; Beauchamp, K. A.; Wang, L.-P.; Simmonett, A. C.; Harrigan, M. P.; Stern, C. D.; Wiewiora, R. P.; Brooks, B. R.; Pande, V. S. OpenMM 7: Rapid development of high performance algorithms for molecular dynamics. PLOS Computational Biology 2017, 13, e1005659.

(59) Shirts, M.; Pande, V. S. Screen savers of the world unite! Science 2000, 290, 1903–1904.

(60) Roe, D. R.; Cheatham, T. E. PTRAJ and CPPTRAJ: Software for Processing and Analysis of Molecular Dynamics Trajectory Data. Journal of Chemical Theory and Computation 2013, 9, 3084–3095.

(61) McGibbon, R. T.; Beauchamp, K. A.; Harrigan, M. P.; Klein, C.; Swails, J. M.; Hernández, C. X.; Schwantes, C. R.; Wang, L.-P.; Lane, T. J.; Pande, V. S. MDTraj: A Modern Open Library for the Analysis of Molecular Dynamics Trajectories. Biophysical Journal 2015, 109, 1528–1532.

(62) Humphrey, W.; Dalke, A.; Schulten, K. VMD: Visual molecular dynamics. Journal of Molecular Graphics 1996, 14, 33–38.

