## Supplementary Methods, Images, Tables and Results for "The Effects of N-linked Glycosylation on SLC6 Transporters"

#### SUPPORTING INFORMATION

This PDF file includes:

Figures S1 to S5

Tables S1 to S3

### Supplementary Figures

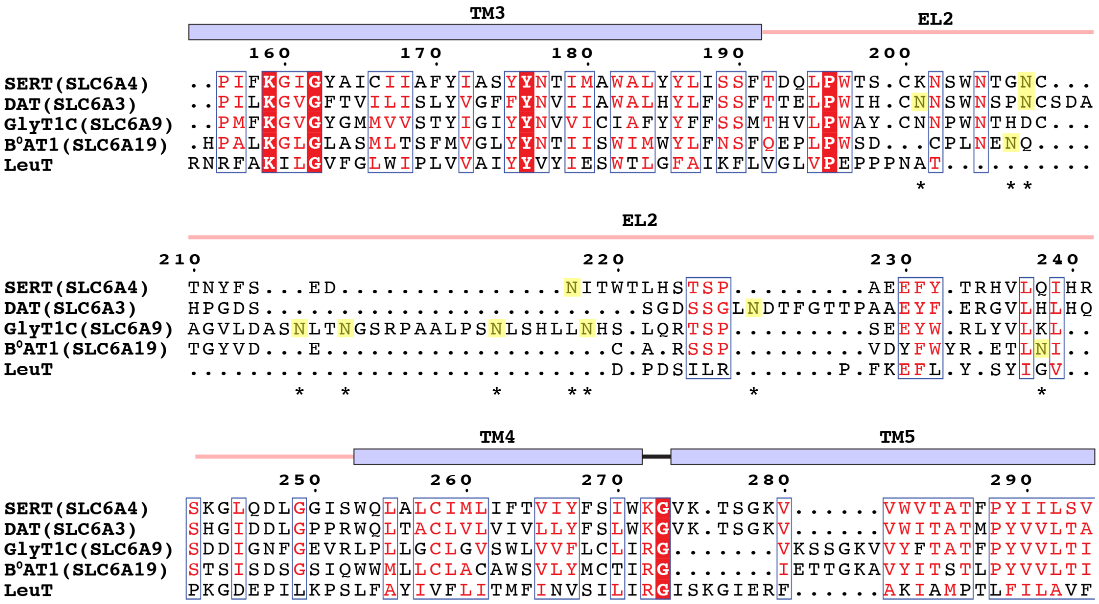


Figure S1: **Multiple sequence alignment of studied SLC6 transporters.** N-linked glycosylation sites are highlighted and indicated by the asterisks (*). The residue number of the MSA corresponds to the SERT sequence.

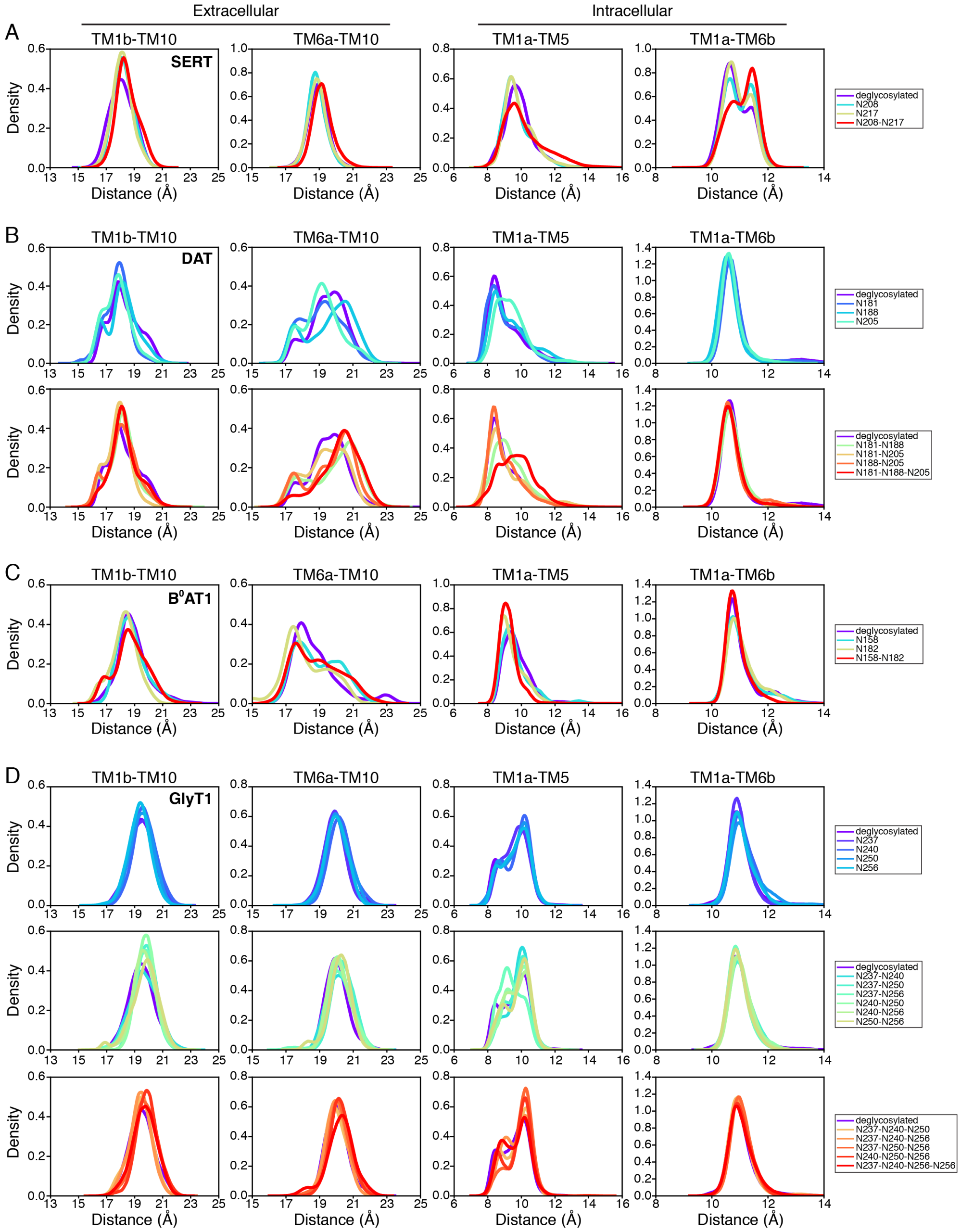


Figure S2: **Distance distribution of gating helices.** The center-of-mass distances between gating helices TM1b-TM10 and TM6a-TM10 on the extracellular side and TM1a-TM5 and TM1a-TM6b on the intracellular side were measured from the 1$\mu$s simulations. The data represent simulations in which a single N-acetylglucosamine glycan was modeled to the specified glycosylated site(s). The glycosylated residues are colored by the transporter's respective legend. Specific residues used for the center-of-mass calculations are detailed in Table S3.

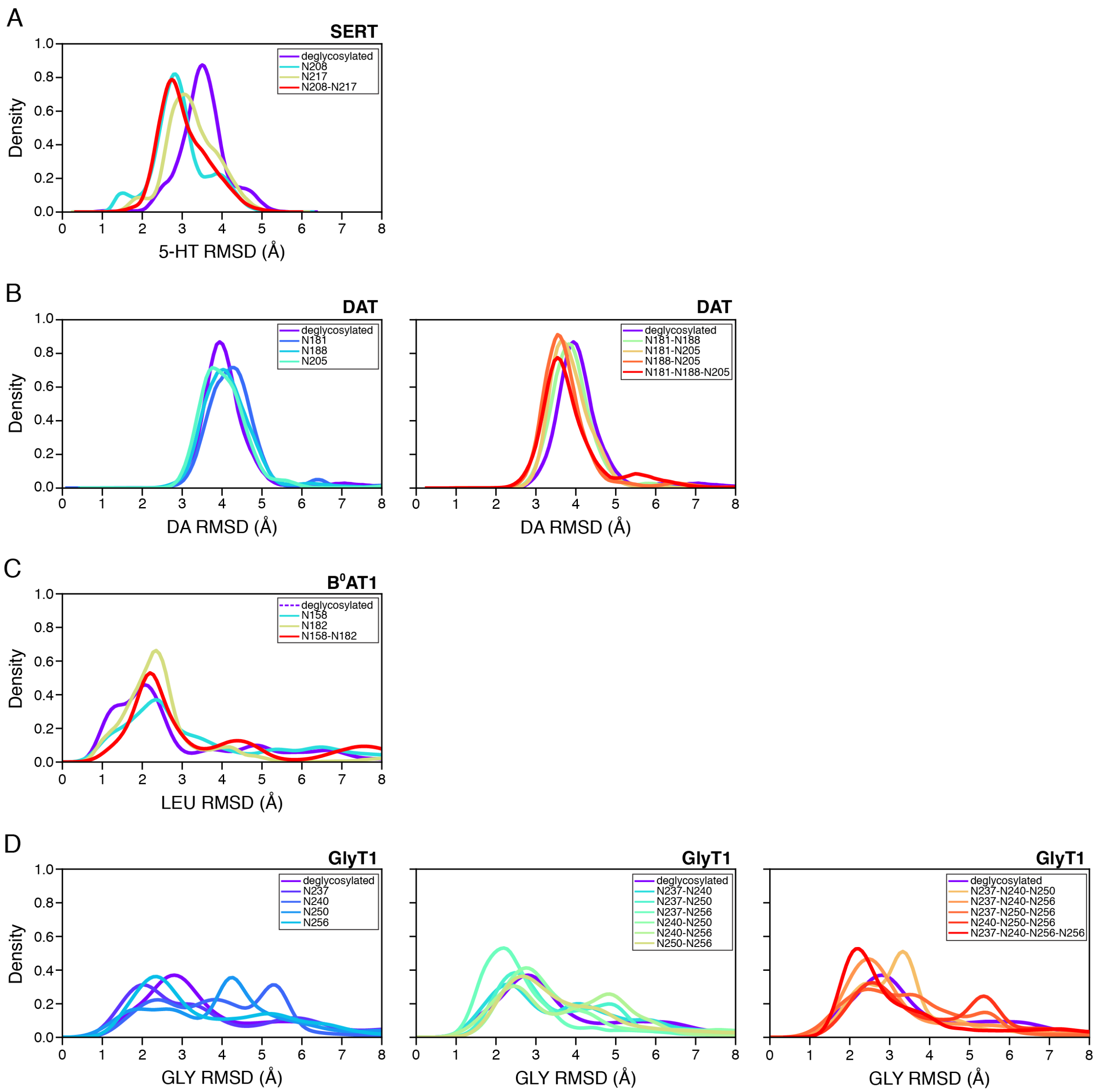


Figure S3: **Stability of the ligand in the orthosteric pocket.** Distribution of the RMSD of the respective ligand with respect to the initial bound pose. The data represent simulations in which a single N-acetylglucosamine glycan was modeled to the specified glycosylated site(s). The glycosylated residues are colored by the transporter's respective legend.

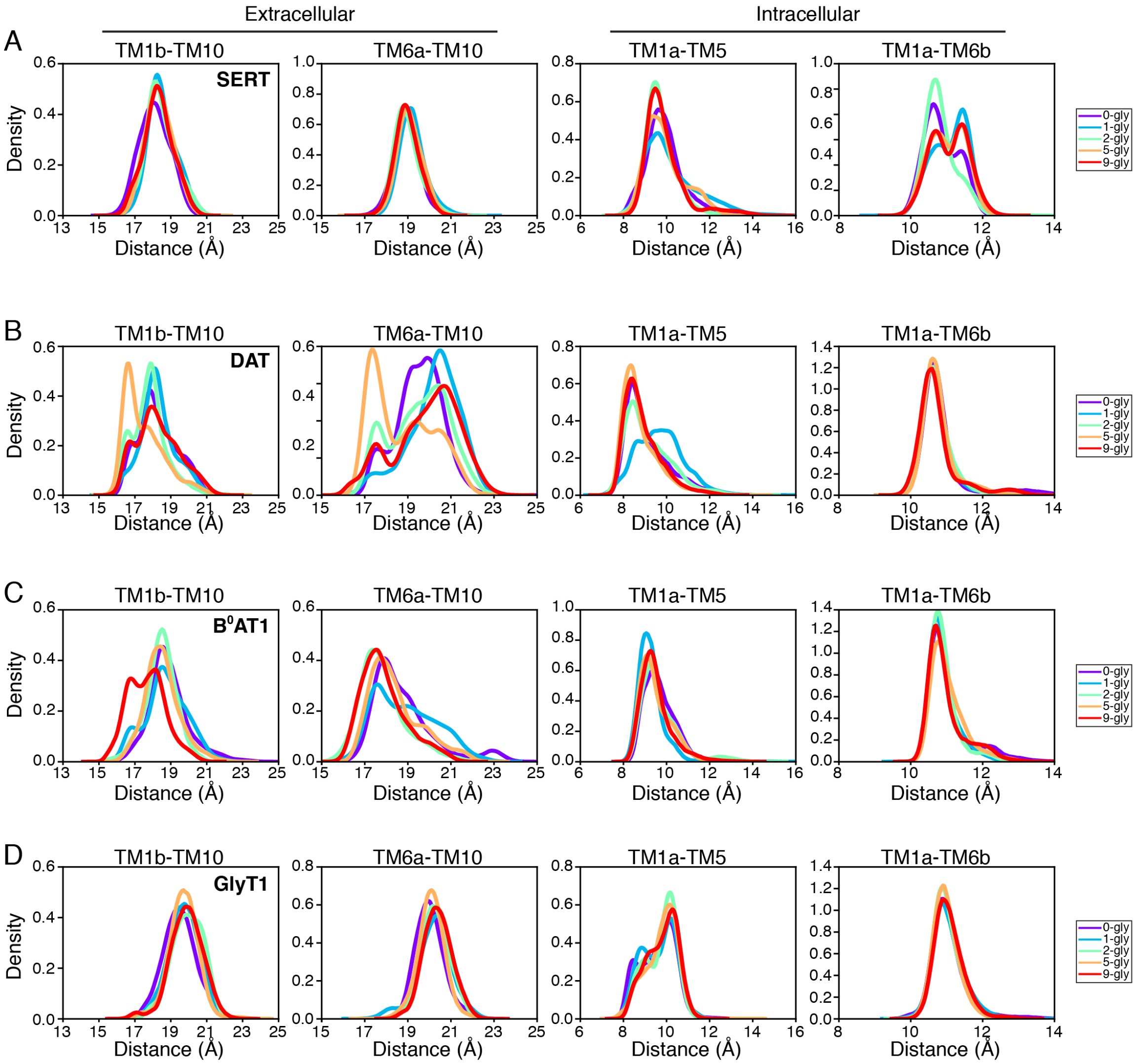


Figure S4: **Distance distribution of gating helices from complex oligoglycan simulations.** The center-of-mass distances between gating helices TM1b-TM10 and TM6a-TM10 on the extracellular side and TM1a-TM5 and TM1a-TM6b on the intracellular side were measured from the 1$\mu$s simulations. The data represent simulations in which an oligoglycan was modeled to the specified glycosylated site(s). See Figure 4 for details of the simulated olgioglycans. The degree of glycosylation are colored by the transporter's respective legend. Specific residues used for the center-of-mass calculations are detailed in Table S3.

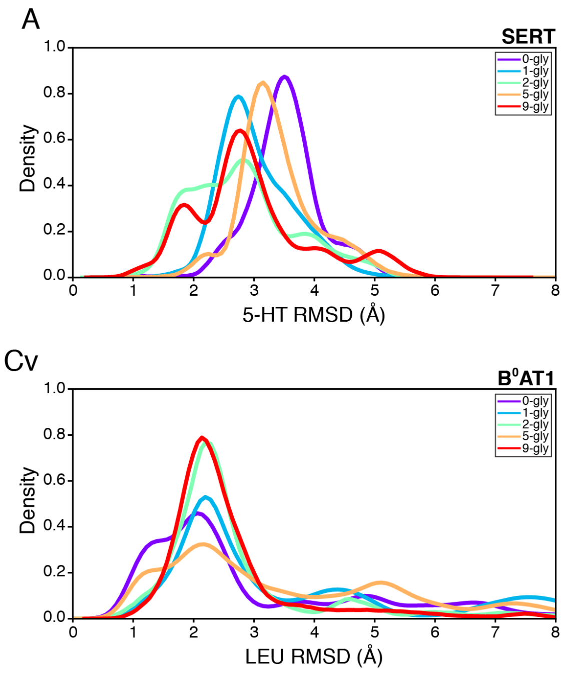


Figure S5: **Stability of the ligand in the orthosteric pocket** Distribution of the RMSD of the respective ligand with respect to the initial bound pose. The data represent simulations in which an oligoglycan was modeled to the specified glycosylated site(s). See Figure 4 for details of the simulated olgioglycans. The degree of glycosylation are colored by the transporter's respective legend.

#### Supplementary Tables

Table S1: **Overview of N-acetylglucosamine-transporter MD simulations.** Simulations were initiated from the outward-facing conformation with respective substrates bound in the orthosteric site and embedded in the described membrane. An N-acetylglucosamine glycan was modeled to the specified glycosylated site(s). Individual trajectories were 1$\mu$s in length.

| **Transporter** | **Substrate bound** | **Membrane** | **Number of glycosylation sites** | **Glycosylated site** | **Number of replicates** |
|---|---|---|---|---|---|
| SERT | 5-HT, Na1, Na2, $Cl^-$ | 2:1 POPC:POPE | 2 | None | 29 |
| | | | | N208 | 30 |
| | | | | N217 | 30 |
| | | | | N208-N217 | 29 |
| DAT | DA, Na1, Na2, $Cl^-$ | 2:1 POPC:POPE | 3 | None | 29 |
| | | | | N181 | 30 |
| | | | | N188 | 29 |
| | | | | N205 | 29 |
| | | | | N181-N188 | 30 |
| | | | | N181-N205 | 30 |
| | | | | N188-N205 | 29 |
| | | | | N181-N188-N205 | 30 |
| $B^0AT1$ | Leu, Na1, Na2 | 3:2:1 POPE:POPC:POPS | 2 | None | 30 |
| | | | | N158 | 29 |
| | | | | N182 | 29 |
| | | | | N158-N182 | 30 |
| GlyT1 | Gly, Na1, Na2, $Cl^-$ | 2:1 POPC:POPE | 4 | None | 30 |
| | | | | N237 | 30 |
| | | | | N240 | 30 |
| | | | | N250 | 29 |
| | | | | N256 | 30 |
| | | | | N237-N240 | 30 |
| | | | | N237-N250 | 29 |
| | | | | N237-N256 | 30 |
| | | | | N240-N250 | 30 |
| | | | | N240-N256 | 30 |
| | | | | N250-N256 | 30 |
| | | | | N237-N240-N250 | 29 |
| | | | | N237-N240-N256 | 30 |
| | | | | N237-N250-N256 | 30 |
| | | | | N240-N250-N256 | 30 |
| | | | | N237-N240-N250-N256 | 30 |

Table S2: **Overview of oligo-N-glycan-transporter MD simulations.** Simulations were initiated from the outward-facing conformation with respective substrates bound in the orthosteric site and embedded in the described membrane. Varying degrees of glycan were modeled to all glycosylation sites. See Figure 4 for details of the olgioglycans. Individual trajectories were 1$\mu$s in length.

| **Transporter** | **Substrate bound** | **Membrane** | **Number of glycosylation sites** | **Number of glycans per glycosylation site** | **Number of replicates** |
|---|---|---|---|---|---|
| SERT | 5-HT, Na1, Na2, Cl- | 2:1 POPC:POPE | 2 | 0 | 29 |
| | | | | 1 | 29 |
| | | | | 2 | 30 |
| | | | | 5 | 30 |
| | | | | 9 | 30 |
| DAT | DA, Na1, Na2, Cl- | 2:1 POPC:POPE | 3 | 0 | 29 |
| | | | | 1 | 30 |
| | | | | 2 | 29 |
| | | | | 5 | 30 |
| | | | | 9 | 30 |
| $B^0AT1$ | Leu, Na1, Na2 | 3:2:1 POPE:POPC:POPS | 2 | 0 | 30 |
| | | | | 1 | 30 |
| | | | | 2 | 30 |
| | | | | 5 | 30 |
| | | | | 9 | 30 |
| GlyT1 | Gly, Na1, Na2, Cl- | 2:1 POPC:POPE | 4 | 0 | 30 |
| | | | | 1 | 30 |
| | | | | 2 | 30 |
| | | | | 5 | 30 |
| | | | | 9 | 30 |

Table S3: Residues considered for center-of-mass calculation of helix distances.

Extracellular helices

| | **TM1b** | **TM6a** | **TM10** |
|---|---|---|---|
| **SERT** | L99-N112 | T323-G338 | G484-T497 |
| **DAT** | A81-G94 | S309-G323 | G468-G481 |
| **$B^0$AT1** | L53-H66 | P266-F280 | S478-L53 |
| **GlyT1** | L121-N134 | A359-G374 | A520-A533 |

Intracellular helices

| | **TM1a** | **TM5** | **TM6b** |
|---|---|---|---|
| **SERT** | K85-A96 | V274-T286 | V343-V102 |
| **DAT** | I67-V78 | K260-M272 | L329-N336 |
| **$B^0$AT1** | K39-C50 | I217-T229 | G286-Y293 |
| **GlyT1** | Q107-A118 | V310-T322 | G379-Y386 |
